# DecoyFinder: Identification of Contaminants in Sets of Homologous RNA Sequences

**DOI:** 10.1101/2024.10.12.618037

**Authors:** Mingyi Zhu, Jeffrey Zuber, Zhen Tan, Gaurav Sharma, David H. Mathews

**Affiliations:** Center for RNA Biology, University of Rochester Medical Center, Rochester, NY, United States; Department of Biochemistry and Biophysics, University of Rochester Medical Center, Rochester, NY, United States; University of Rochester, Department of Electrical and Computer Engineering, Rochester, NY, United States; University of Rochester, Department of Computer Science, Rochester, NY, United States

## Abstract

**Motivation:** RNA structure is essential for the function of many non-coding RNAs. Using multiple homologous sequences, which share structure and function, secondary structure can be predicted with much higher accuracy than with a single sequence. It can be difficult, however, to establish a set of homologous sequences when their structure is not yet known. We developed a method to identify sequences in a set of putative homologs that are in fact non-homologs.

**Results:** Previously, we developed TurboFold to estimate conserved structure using multiple, unaligned RNA homologs. Here, we report that the positive predictive value of TurboFold is significantly reduced by the presence of contamination by non-homologous sequences, although the reduction is less than 1%. We developed a method called DecoyFinder, which applies machine learning trained with features determined by TurboFold, to detect sequences that are not homologous with the other sequences in the set. This method can identify approximately 45% of non-homologous sequences, at a rate of 5% misidentification of true homologous sequences.

**Availability:** DecoyFinder and TurboFold are incorporated in RNAstructure, which is provided for free and open source under the GPL V2 license. It can be downloaded at http://rna.urmc.rochester.edu/RNAstructure.html

## 1 Introduction

RNA plays an essential role in all domains of life. RNA was once thought to only have roles in the expression of proteins, but we now understand that RNA also functions in diverse roles, including catalysis and gene regulation, much like proteins (Doudna and Cech, 2002; Serganov and Nudler, 2013). The RNAs that function other than by encoding proteins are called non-coding RNA (ncRNA) (Eddy, 2001). The structures for many ncRNAs are critical for their function (Cech, 2002; Somarowthu, et al., 2015; Vandivier, et al., 2016).

RNAs tend to fold hierarchically by base pairing and then by forming additional contacts (Tinoco and Bustamante, 1999). RNA secondary structure is the set of canonical base pairs (i.e., A-U, C-G and G-U). The base pairs are in A-form helices, and the residues not in canonical base pairs are in loops, which include bulge loops, multibranch loops, internal loops, and exterior loops (Smit, et al., 2009). The secondary structure is conserved across orthologous sequences, a.k.a. RNA families, and the secondary structure can be used to understand function.

With the development of next generation sequencing, it is possible to obtain complete genomes and transcriptomes (Behjati and Tarpey, 2013; Ben Khedher, et al., 2022). Algorithms can be used to scan these data to identify new instances of established RNA families, once the conserved secondary structure is established (Nawrocki and Eddy, 2013). Databases have been developed to catalog the sequences that are assigned to specific RNA families (Hoeppner, et al., 2014).

RNA secondary structure can be predicted for ncRNAs that are not part of established families. Free energy minimization is a popular method for predicting RNA secondary structure (Andronescu, et al., 2014; Hofacker, 2014; Seetin and Mathews, 2012). The average accuracy of single sequence secondary structure is approximately 70% for sequences shorter than 700 nucleotides (Hofacker, 2014; Mathews, et al., 2004). Because RNA structures are conserved across evolution, in spite of the mutations in the sequences (Brown, et al., 2009; Rivas, et al., 2017), predicting the conserved structure using a set of homologs can improve the accuracy of the RNA secondary structure prediction. We developed TurboFold, which takes unaligned homologs as input and estimates both the sequence alignment and the base pairing probabilities of the sequences. TurboFold iteratively refines each, using the alignment to facilitate the prediction of consensus base pairs and the predicted base pairs to guide the alignment. TurboFold improves prediction sensitivity compared to single sequence predictions and has an average structure prediction sensitivity of approximately 80% (Tan, et al., 2017). Other methods have also been developed to estimate conserved RNA structures, and reviews summarize these approaches (Asai and Hamada, 2014; Havgaard and Gorodkin, 2014; Mathews, et al., 2010).

To determine a conserved structure, a set of homologous sequences is needed. Sequence alone is generally not sufficient to identify homologs for structured RNAs. Unlike proteins, where amino acids have distinct biophysical characteristics that limit substitutions across evolution, RNA bases can all serve equivalent roles through covariations that conserve structure (Michel and Westhof, 1990; Woese and Pace, 1993). To find homologs, a computational tool exists for finding homologs from a single instance (Klein and Eddy, 2003). Other computational tools automate the discovery and alignment of sets of homologs (Eggenhofer, et al., 2016; Zhang, et al., 2022). Sometimes, however, homolog discovery requires manual effort using biological insight or synteny across genomes (Jakubczak, et al., 1991; Kiontke, et al., 2019). Each of these methods are prone to false discovery.

The discovery of false homologs is an ongoing concern during the establishment of new RNA families. The nonhomologous sequences represent contaminant sequences, and we refer to them as decoys. Here, we report the extent to which structure prediction accuracy is affected when decoys are present in a group of homologous sequences. The presence of decoy sequences, constituting 20% of all sequences, reduces structure prediction accuracy of the homologous sequences by TurboFold, with mean sensitivity (the fraction of pairs in the accepted structure that are predicted) reduced by 0.15% and mean positive predictive value (PPV; the fraction of predicted pairs in the accepted structure) reduced by 0.67% across RNA families. We also discovered that the data generated by TurboFold calculations can be used to detect decoy sequences. We present an algorithm called DecoyFinder (Figure 1), which uses data calculated by TurboFold and a machine learning model to identify the decoy sequences from a group of putative homologous sequences. DecoyFinder can be tuned to have higher sensitivity or specificity for decoy discovery. At a threshold that identifies about 45% of the decoy sequences, less than 5% of the homologous sequences are misidentified. DecoyFinder provides a new tool for curation of homologous sequences in the stages before a family is firmly established.

**Figure 1.**
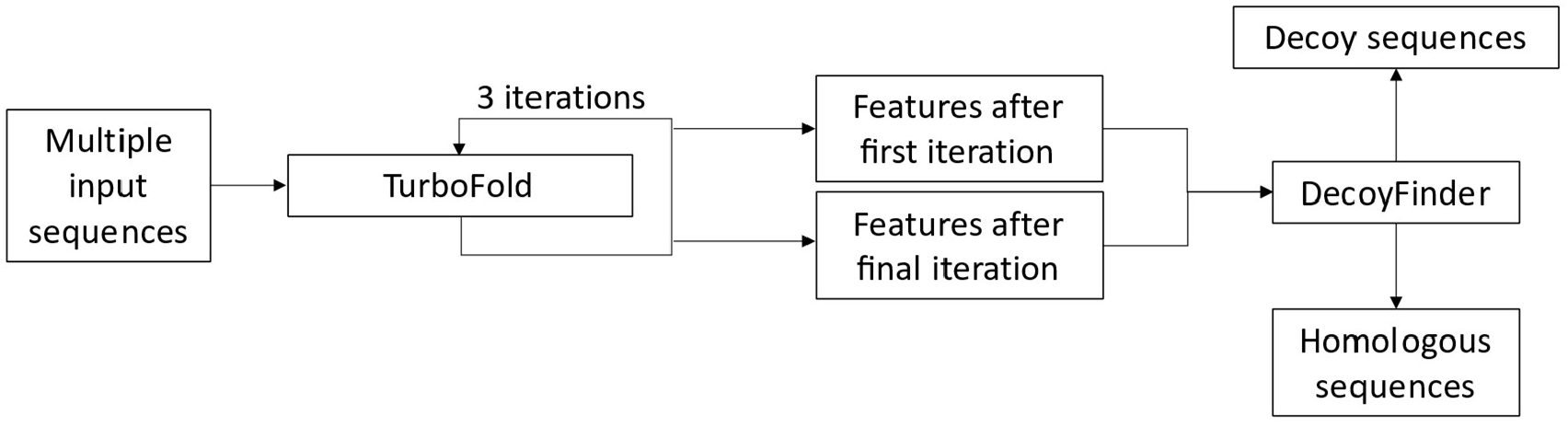
Overview of DecoyFinder. TurboFold takes a set of putative homologous sequences. TurboFold runs 3 iterations to refine the estimated alignment and structures. Feature data after the first and final iterations from TurboFold are used by DecoyFinder, and, according to the machine learning model, DecoyFinder classifies each sequence as homologous to the sequence set or as a decoy to the set.

## 2 Methods

### 2.1 TurboFold and features it determines for decoy classification

TurboFold is an algorithm we developed to predict conserved RNA secondary structure that utilizes iterative refinement of sequence alignment and partition function calculations of base pairing probabilities (Harmanci, et al., 2011; Tan, et al., 2017). TurboFold is a component of the RNAstructure package, and calculations reported here used RNAstructure version 6.2.

In TurboFold, all pairs of sequences are initially aligned using a Hidden Markov Model (HMM) (Harmanci, et al., 2007). The base pairing probabilities of each sequence are also initially estimated by partition function calculations for each sequence (Mathews, 2004; McCaskill, 1990). We call these pairing estimates the “intrinsic information.” Next, a proclivity of base pairing for each pair in each sequence is estimated using base pairing probabilities predicted for other sequences and the alignment of the sequences, and we call this “extrinsic information.” The extrinsic information is used to refine base pairing probabilities in subsequent iterations of the partition function calculations, so that the predicted base pairing probabilities reach a consensus. Also using the estimated pair probabilities, match scores (defined below) are calculated to quantify the similarity in structure between aligned nucleotide positions in two sequences (Hofacker, et al., 2004). The match scores are used in subsequent iterations of the alignment HMM to refine the alignments. The match score is higher when aligned nucleotides are both upstream paired, downstream paired or unpaired. By using the match score in the HMM, the nucleotides with the same pairing status are more likely to align. After multiple iterations of the alignment and base pair probability estimates (three by default), a multiple sequence alignment (using a probability consistency transformation) is generated and maximum expected accuracy structures for each sequence are predicted (Do, et al., 2005; Lu, et al., 2009).

#### Match score

The match scores are calculated in TurboFold using the base pairing probabilities (Hofacker, et al., 2004). Generally, aligned nucleotide positions in homologous sequences have conserved base pairing status, which can be upstream paired (>), downstream paired (<), or unpaired (o). For the nucleotides *i* and *k* in sequences *m* and *n*, respectively, the match score is formulated as:

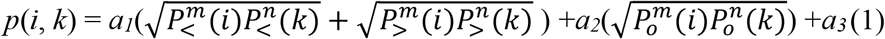

where 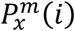 represents the probability of nucleotide *i* in sequence *m* being in pairing status *x* (paired downstream, >; paired upstream, <; or unpaired, *0*). The 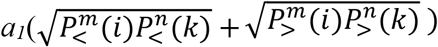 term accounts for conserved pairing (upstream or downstream). 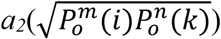 accounts for the case where *i* and *k* are both unpaired. *a*_*1*_, *a*_*2*_, and *a*_*3*_ are weight parameters, optimized previously by a grid search for parameters that maximized prediction accuracy: *a*_*1*_ = 1.0, *a*_*2*_ = 0.8, and *a*_*3*_ = 0.5 (Tan, et al., 2017).

#### Pairing and Unpairing Score (PS and US)

We found that alignments of decoys generated by TurboFold do not conserve the pairing status as well as homologs. From an estimated alignment, we calculate a pairing score (*PS*^*m*^(*i*)), summarizing upstream and downstream pairing, and unpairing score (*US*^*m*^(*i*)) of alignment column (*i*). Both pairing score (*PS*) and unpairing score (*US*) derive from the match score, and are formulated as:

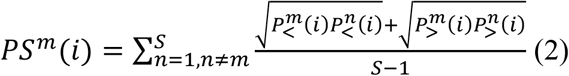

and

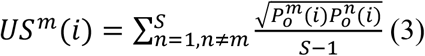

where *n* iterates across the sequences and *S* is the number of sequences in the alignment. *PS*^*m*^(*i*) and *US*^*m*^(*i*) are calculated for each column, *i*, for each sequence *m*. If position *i* in sequence *m* is a gap, then the probability of pairing and unpairing is zero, and column *i* has *PS*^*m*^(*i*) and *US*^*m*^(*i*) equal to zero. For a given sequence, *m*, we generate a two-dimensional array of counts for *PS*^*m*^(*i*) and *US*^*m*^(*i*) across *i*, i.e., a two-dimensional histogram. The x-axis (*PS*) and y-axis (*US*) of the array represent probabilities spanning the range from 0 to 1 and discretized using a bin size of 1/30. Because *PS*^*m*^(*i*) + *US*^*m*^(*i*) ≤ 1.0, only the lower triangular elements of the array need to be stored. Therefore, the array generated is triangular with 465 elements, truncated from the 30×30 two-dimensional array by removing all elements above the diagonal that connects (*PS*^*m*^(*i*) *=* 1, *US*^*m*^(*i*) = 0) and (*PS*^*m*^(*i*) *=* 0, *US*^*m*^(*i*) = 1). The values in the array are normalized by the total counts in the array. The interval of scale was optimized to maximize machine learning accuracy on the training set as described below. This truncation was achieved.

#### Kullback-Leibler divergence score (KL score)

We found that using the full matrix of 465 elements in the *PS*(*i*) and *US*(*i*) array did not allow the machine learning to integrate other sources of data. To reduce the *PS*^*m*^(*i*) and *US*^*m*^(*i*) matrix to a single metric, the Kullback-Leibler divergence (KL divergence), a measure of difference between probability distributions (Kullback and Leibler, 1951), was used. For the pairing and unpairing score of sequence *m* with sequences *n, n≠m*, the *KL score* is formulated as:

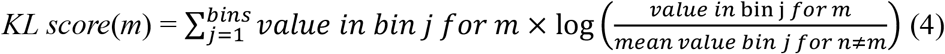

where j iterates across all the bins in the array. To avoid problems with zero population for *m* or for zero mean population, we use pseudocounts (Wilson, 1927), where the value of each bin is incremented by (1/total bin counts) prior to the computation in (4).

### 2.2 Additional features

#### Z-score

The Z-score of folding free energy change represents how different the folding free energy (ΔG°) of sequence *m* is from the other sequences *n* (*n≠m*) using the calculation:

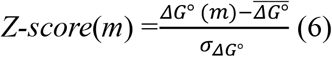

where the mean free energy change, 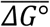, and the standard deviation of the free energy change, *σ*_*ΔG*°_, are calculated using all sequences except *m*. The folding free energy change is calculated by the efn2 program in RNAstructure using the Turner 2004 nearest neighbor rules (Reuter and Mathews, 2010; Turner and Mathews, 2010).

#### Mean Sequence Shannon entropy

This feature reflects how well a sequence aligns with other sequences in the group, and is formulated as:

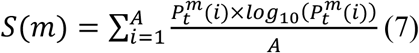

where *i* is the position in the alignment of length *A*, t is the nucleotide identity at position *i* for sequence m, 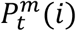 is the probability of nucleic acid base identity *t*, where gaps are included as an identity along with nucleotides, for position *i*.

#### Structural Shannon entropy

The structural Shannon entropy provides a measure of well-definedness for the formation of a single structure (Huynen, et al., 1997). It is calculated as:

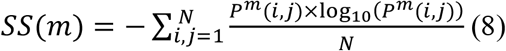

where *P*^*m*^(*i, j*) is the probability that nucleotides *i* and *j* are base paired in sequence *m* and *N* is the length of sequence *m*. For a feature in decoy discovery, we use the difference between *SS*(*m*) after the 0^th^ iteration of TurboFold (i.e. the single sequence estimates of Shannon Entropy) and after all TurboFold iterations. We hypothesize the decoy sequences have greater change because TurboFold forces them to fold into a structure that is closer to the homologous sequences.

### 2.3 Shuffled decoy sequences

To train and test our method, we needed decoy sequences that would align as well as homologous sequences to the set of homologous sequences. To accomplish this, we shuffled a fraction of positions of a homologous sequence to generate a decoy, keeping the unshuffled nucleotides constant. The fraction of positions to keep constant was determined by RNA family and is the mean sequence identity within the family from which the homologs are derived. Therefore, the shuffled sequence (decoy) will have a minimum identity equal to the mean pairwise nucleotide identity for the family.

The positions to shuffle are chosen at random and then subjected to the usual shuffling algorithm.

### 2.4 Machine learning and training sets

The Adaptive Boosting (AdaBoost) Classifier in the Scikit-learn package (Pedregosa, et al., 2011) was trained with 2100 training groups, drawn from the Rfam families 5S rRNA, internal ribosome entry site (IRES), malK-I RNA, RAGATH-18 RNA, RNase P RNA, skipping rope RNA, tRNA and U3 RNA (Kalvari, et al., 2020). Each set has 5 to 20 homologous sequences (taken from the family) and 1 to 3 decoy sequences in each group that are from other training families (taken from other families randomly) or are shuffled decoy sequences. 600 of these groups were used for each of 1, 2, and 3 decoy sequences with the homologs. There are an additional 300 training groups with 5 to 20 homologous sequences from the same family as training family list above, but no decoy sequences (Supplementary Table 1). We also tested linear regression, ridge regression, support vector machines (SVM), support vector classifier (SVC), Naive Bayes, Gaussian Naive Bayes, decision trees, multi-layer perceptron (MLP), random forest, and the KNeighborsClassifier. By comparing the area under the ROC curve (AUC) of different methods, we found that Adaptive Boosting performed the best, so it was selected to be the machine learning model for DecoyFinder.

#### Parameter optimization

We used a grid search with the Scikit-learn package (Pedregosa, et al., 2011), using the AdaBoost Classifier as the estimator and the area under the ROC curve as scoring metric, to optimize the histogram bin size for the *PS*-*US* histogram array. The parameters, the maximum number of estimators (from 20 to 20000), learning rate (0.5 to 20) and algorithm (‘SAMME’ and ‘SAMME.R’) of the AdaBoost Classifier were optimized by grid search using area under the ROC curve as scoring metric from Scikit-learn.

### 2.5 Testing sets

700 testing groups (drawn from RNA families pemK RNA motif, small nucleolar RNA SNORD116, U4 spliceosomal RNA, U5 spliceosomal RNA or Y RNA) were generated in the same manner as the training sets in order to validate the trained model and contains no overlap of RNA families in the training and testing sets (Lu, et al., 2009; Rivas, et al., 2012; Szikszai, et al., 2022). The selection of homologous and decoy families was random. The homologous sequences in all sets are randomly selected, ranging from 5 to 20. For decoys drawn from other families, there are 100 test sets for each of one decoy, two decoys, and three decoys in the set. For shuffled decoys, there are also 100 sets for each of one decoy, two decoys, and three decoys in the set. There are an additional 100 test sets that have 5 to 20 homologous sequences from the same family as testing family list above, and no decoy sequences (Supplementary Table 2).

### 2.6 Infernal calculations

Infernal (version 1.1.4, Released on December 9, 2020) was tested for its ability to determine whether a sequence belongs to a homologous family or to other families (Nawrocki and Eddy, 2013). We used the 700 test groups we assembled for testing DecoyFinder (Supplementary Table 2). Each test group is made up of 5 to 20 homologous sequences and 1 to 3 decoy sequences selected from other testing families or shuffled. The Infernal-trained models were downloaded from Rfam for each RNA family. Each sequence was categorized using the default threshold.

### 2.7 Single sequence structure prediction

Single sequence structure prediction was performed with the MaxExpect program in RNAstructure, which predicts maximum expected accuracy structures from predicted base pairing probabilities (Do, et al., 2005; Lu, et al., 2009).

### 2.8 Accuracy of secondary structure prediction

The accuracy of secondary structure prediction as compared to known structures was calculated by the scorer program in the RNAstrucure package. Scorer compares a predicted structure to an accepted structure and returns the sensitivity and positive predictive value (PPV) (Mathews, 2019). To test for significance for changes in these accuracy metrics, a two tailed, paired t test was performed with the Python package SciPy (version 1.5.4) (Virtanen, et al., 2020; Xu, et al., 2011).

## 3 Results

### 3.1 Development of Decoy Sequences

In this study, we used two types of decoys to train and test our machine learning method. The first type of decoy is a sequence that is drawn from a different Rfam family, which therefore does not share the structure with the homologs. The second type of decoy is a sequence that is from the same family with partial sequence shuffling.

We generate family-specific shuffled decoy sequences by randomly choosing a homolog and then shuffling a fraction of nucleotides, with the remaining nucleotides fixed in position and identity. The fraction of fixed positions is the mean sequence identity within the RNA family. These shuffled decoy sequences have a sequence identity that would therefore make them appear like a member of the RNA family because they have the proper length for a member of the family and should align as well with the homologous sequences as a true homolog.

We found that the shuffled decoy sequences have a different predicted secondary structure than the original sequence, in spite of the relatively high identity to the family. We tested whether shuffled decoy sequences have different predicted pairs than the original sequences using 5S rRNA, 16S rRNA, RNase P RNA, telomerase RNA, tRNA and tmRNA sequences drawn from the RNAStralign database (Brown, 1999; Juhling, et al., 2009; Podlevsky, et al., 2008; RNAcentral Consortium, 2021; Szymanski, et al., 2016; Tan, et al., 2017; Zwieb and Wower, 2000). The secondary structures of the original sequence and shuffled sequence were both predicted using the single sequence and compared to the known structure in the RNAStralign database. The average change in sensitivity between the unshuffled and shuffled sequences was -55.98% and the change in PPV was -49.14% (Supplementary Figure 1). Most base pairs predicted for shuffled sequences are therefore different from those predicted for the original sequence. This demonstrates that shuffled decoy sequences likely form a different structure than the original sequences.

### 3.2 Infernal identifies decoys, but with less accuracy for shuffled decoys

We additionally tested whether Infernal, a method that finds sequence homologs in genome scans, also recognizes that the decoys are not homologous sequences (Nawrocki and Eddy, 2013). Infernal uses a covariance model for each family, and the models were trained using Rfam seed alignments. We used the Rfam seed alignments to generate the shuffled decoys in this test (Kalvari, et al., 2020). We used a total of 700 test groups, including sequences from the pemK RNA motif, small nucleolar RNA SNORD116, U4 spliceosomal RNA, U5 spliceosomal RNA and Y RNA. Each test group is made up of 5 to 20 homologous sequences and 1 to 3 shuffled decoy sequences or decoy sequences (Supplementary Table 2).

We used a receiver operating characteristic (ROC) curve to assess performance (Fig. 2). As expected, the results show that Infernal can correctly identify almost all the decoy sequences that come from alternative families with almost no false prediction of homologous sequences. On the other hand, Infernal has a harder time distinguishing the shuffled decoys from the homologs. For example, at the point that Infernal correctly classifies 95% of the homologs, it can only identify 78.24% shuffled decoy sequence correctly as decoys. This suggests that shuffled decoy sequences are more difficult to distinguish from homologous sequences than sequences derived from other families.

**Figure 2.**
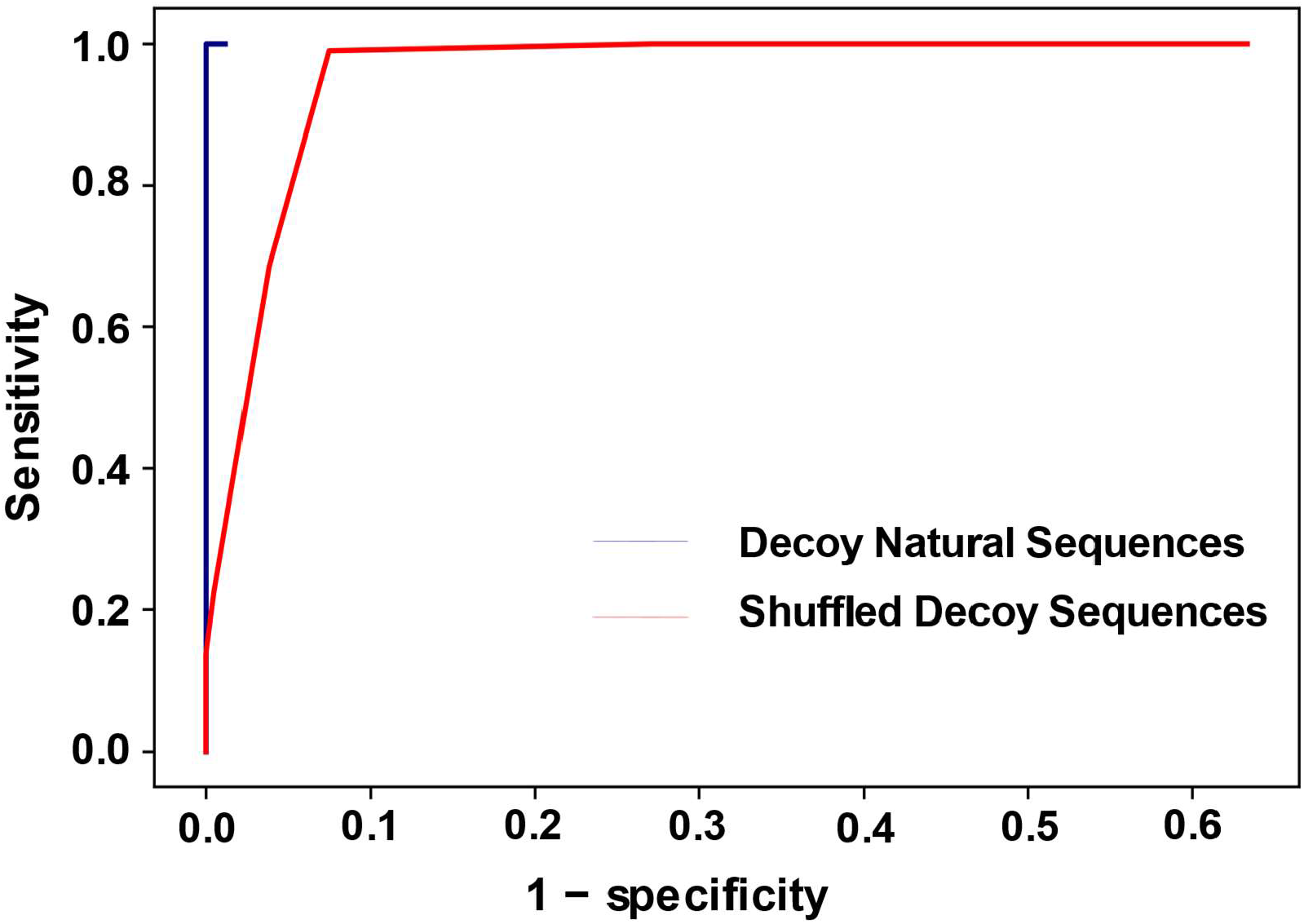
The ROC curve for the Infernal prediction of decoys. The blue line represents the decoys that are natural sequences, drawn from alternative families and the red line represents the decoys that are shuffled from homologous sequences. The sensitivity is the fraction of decoys identified as decoys. The specificity is the fraction of homologs identified as homologs, i.e. a member of the Rfam family.

### 3.3 The accuracy of TurboFold is reduced with both types of decoy sequences

We tested whether the accuracy of TurboFold is adversely affected by having decoy sequences included with a set of homologous sequences in the input. We tested this with both types of decoy sequences, decoy sequences from other RNA families and shuffled decoys. Ideally, TurboFold would be robust to having decoy sequences contaminating the set of input sequences.

We used RNAStralign, which includes alignments and structures of 5S rRNA, 16S rRNA, RNase P RNA, telomerase RNA, tRNA and tmRNA, as the database to test the effect of decoys on the accuracy of homologous sequences secondary structure prediction (Tan, et al., 2017). Each test group is made up of 5 homologous sequences and 1 decoy. All sequences were randomly selected (Supplementary Table 3, Supplementary Table 4). In this test, we have an additional control set for each group of sequences. This set excludes decoy sequences and retains only the homologous sequences. This allows us to determine the effect of decoy sequences on accuracy (Supplementary Table 3).

When RNase P RNAs are mixed with decoys derived from other rfam families, we find that the sensitivity of structure prediction in the homologs decreased 0.22% (p = 0.52) and the positive predictive value reduced by a statistically significant 1.17% (p = 0.01) on average (Figure 3A). The sensitivity of structure prediction in the decoys reduced by a statistically significant 14.37% (p < 0.01) and the PPV reduced by a statistically significant 20.89% (p < 0.01) on average. The RNase P RNA family, however, has an increased sensitivity and a reduced PPV on average.

**Figure 3.**
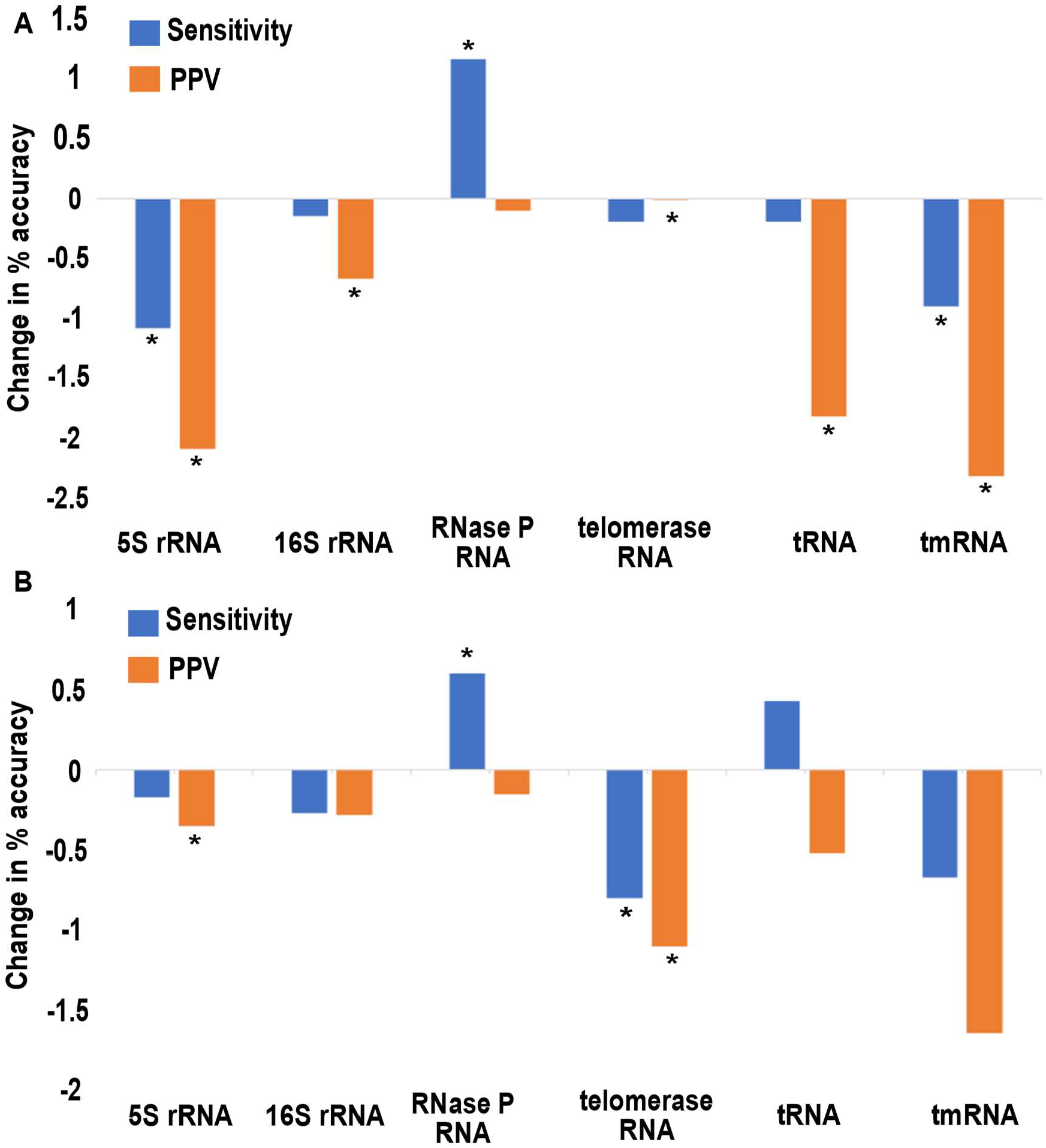
Prediction accuracy difference between TurboFold-predicted MEA structure of sequences with and without decoy sequences sequences included in calculation. (A) The test group with decoys draw from other Rfam families. TurboFold calculation is an average over calculations that include 400 homologs of each RNA family and 100 decoy sequences of each RNA family. (B) The shuffled decoy sequences test group. The TurboFold calculation totally includes 400 homologous of each RNA families and 100 shuffled sequences of each RNA families. Accuracies are calculated for only the homologous sequences, therefore not including decoys. Sensitivity and PPV difference were calculated by comparing TurboFold result with the known structure. The * represents significant changes in accuracies (p < .05).

Next, we tested the effect of including shuffled decoy sequences on prediction accuracy. The decoys are those used for the single sequence benchmark with shuffled decoy sequences (Section 3.1). We observe the mean sensitivity is reduced by 0.15% (p = 0.55) and mean PPV is reduced in a statistically significant manner by 0.67% (p =0.04) across RNA families (Figure 3B). These results suggest that shuffled decoy sequences have even less effect on the prediction accuracy compared to the effect of decoy sequences from other families.

We checked the degree to which the predicted structure for the shuffled decoy matches the structure of the original sequence. Using the structure of the original sequence as the expected target structure, we calculated the accuracy of prediction. The results show that the shuffled decoys can be folded to structures similar to the original homologs when included in a TurboFold calculation. The structure prediction sensitivity significantly increased by 12.4% (p = 0.016) and PPV is significantly increased by 19.3% (p = 0.014) for predictions by TurboFold as compared to a single sequence (Figure 4). These results suggest that shuffled decoy sequences can be difficult to identify and a machine learning method to identify them would be valuable.

**Figure 4.**
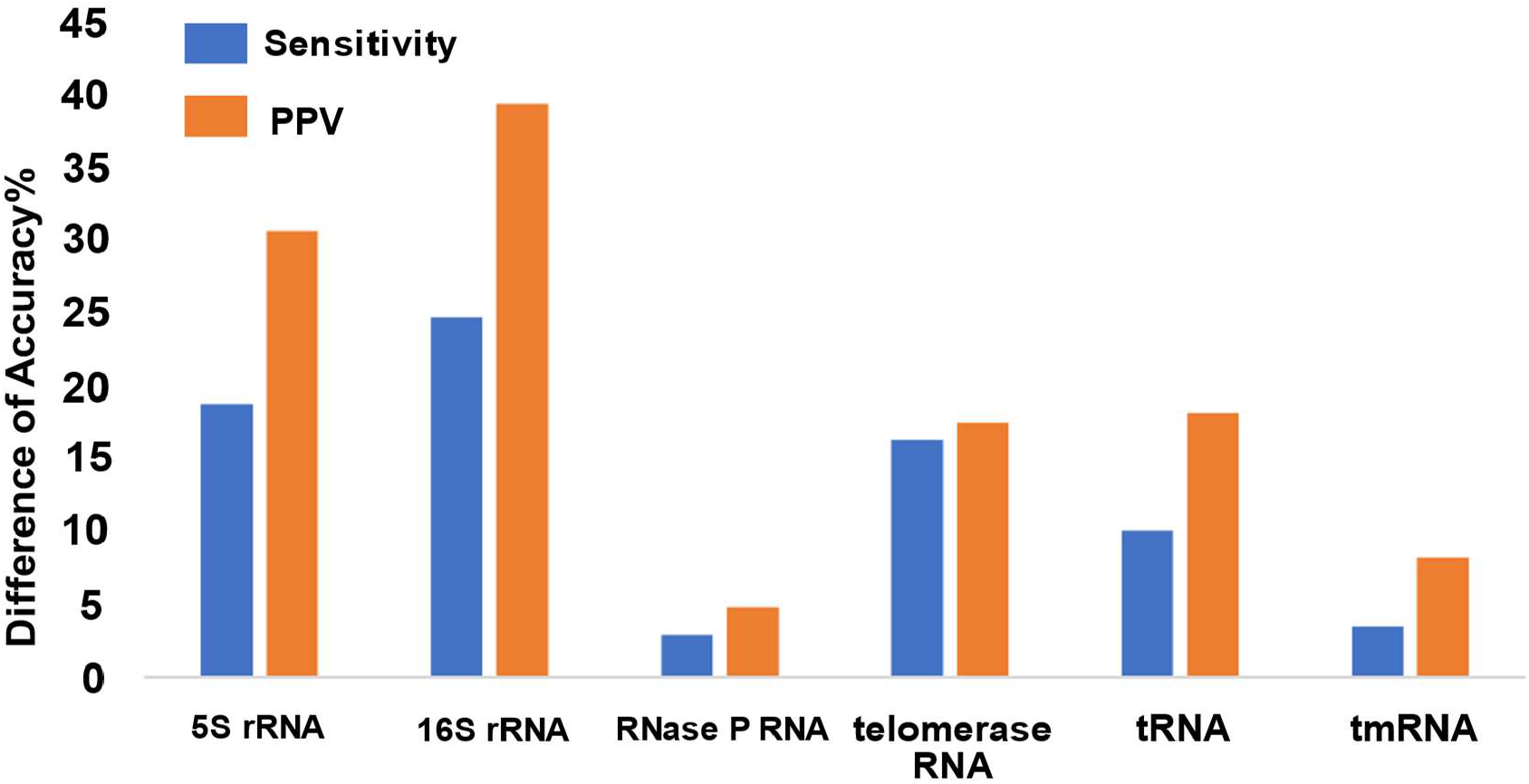
Prediction accuracy difference between single sequence structure prediction and TurboFold structure prediction for shuffled decoys. The TurboFold calculation includes 400 homologous sequences of each RNA families and 100 shuffled sequences of each RNA families. Sensitivity and PPV difference were calculated for MEA structure prediction. Predicted structures are compared to the known structure for the original, unshuffled sequence. The positive values indicate that the shuffled sequence accommodates the conserved structure because the TurboFold-predicted structure is closer to the known structure than the single sequence structure prediction.

To understand why the decoys had such a change in structure prediction accuracy and the TurboFold predictions of decoys are more similar to the structure of the starting sequence, we looked at specific examples. One example is with 9 homologous 5S rRNAs and one shuffled decoy, shuffled from another 5S rRNA (Figure 5). The predicted structure of the 5S shuffled decoy has 41 different base pairs in the TurboFold calculation as compared to single sequence predicted structure (Figure 5B). Using single sequence secondary structure prediction of the shuffled decoy, a substantially different prediction is made (Figure 5C). Apparently, TurboFold is predicting a structure for the decoy that mimics the structure of the homologs. That is because the structure prediction of decoy sequences is guided by the extrinsic information from other sequences, leading to a consensus structure that is more like the conserved structure for the family.

**Figure 5.**
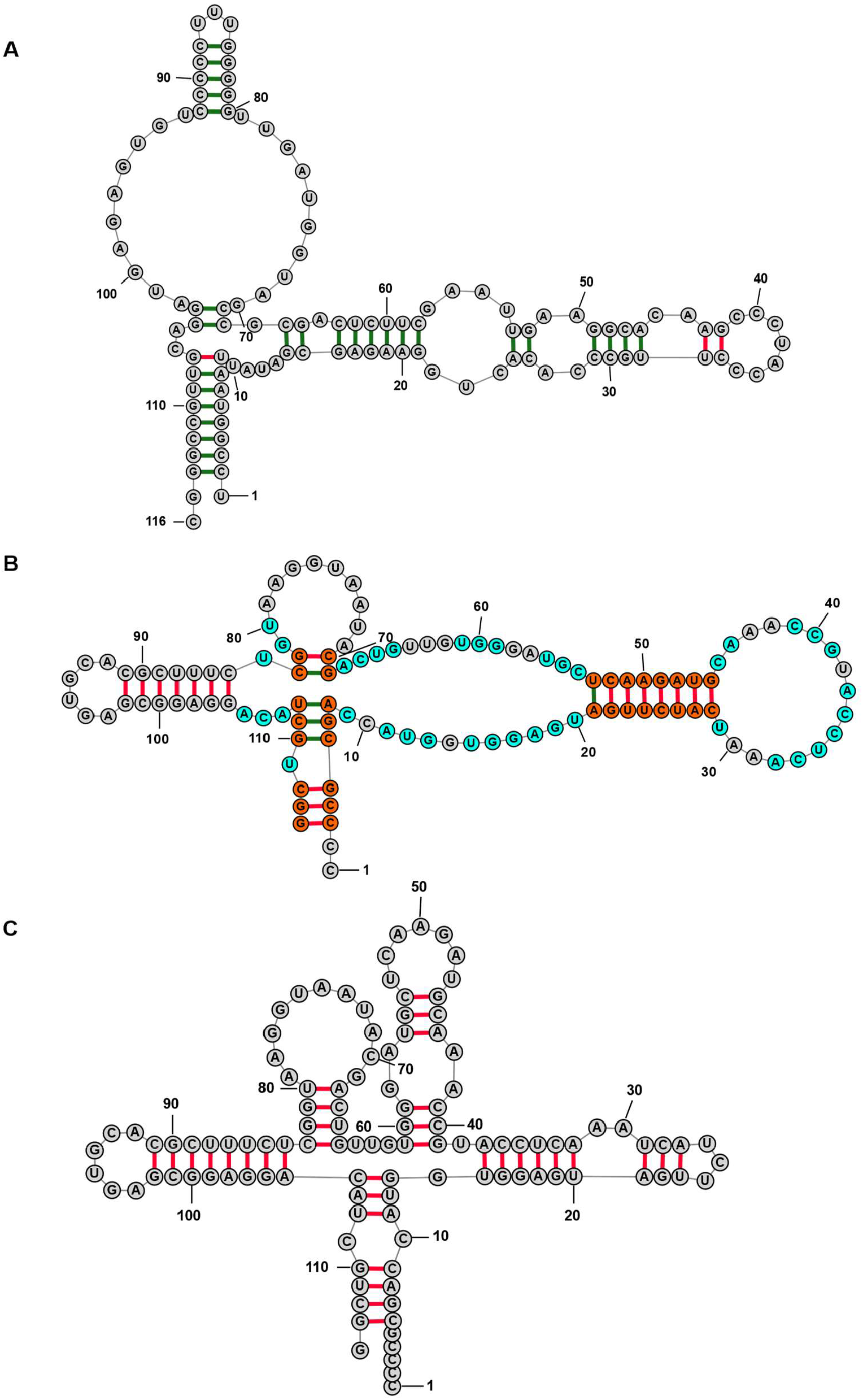
5S rRNA structure prediction using TurboFold including one shuffled decoy. The calculation used nine 5S rRNA and one shuffled decoy. (A) The structure predicted for a 5S rRNA sequence (*Bacillus ginsengihumi*; 5S rRNA Database accession #B05310). (B) The predicted structure for the shuffled decoy (*Haemophilus sputorum*; 5S rRNA Database accession #B03224). (C) The 5S shuffled decoy rRNA (B03224) structure predicted using maximum expected accuracy with a single sequence (Lu, et al., 2009) In panel (B), the orange-colored nucleotides are those in a different base pair compared to panel (C) prediction, and the cyan-colored nucleotides are those base pairs that are not in the panel (C) prediction. In all panels, predicted pairs that occur in the known structure are labeled green and the pairs that are not in the known structure are labeled red.

### 3.4 Decoy sequences stand out in TurboFold calculations

To achieve the most accurate secondary structure prediction and to properly annotate sequences, we want to identify and remove decoy sequences from sets of putative homologs. Therefore, we looked for features that separate decoy sequences from homologs.

One feature that distinguishes decoy sequences is the match score. In TurboFold, the match score, which reflects the structural similarity between two aligned nucleotides from different sequences, is used to refine the pairwise alignments. The match score utilizes the estimated upstream base pairing probability, downstream base pairing probability and unpaired probability to represent the status of aligned nucleotide positions (Eq. 1) (Hofacker, et al., 2004; Tan, et al., 2017). Aligned nucleotides should generally have the same pairing status because homologous sequences have conserved structures, but decoy sequences will not share the conserved structure (Brown, et al., 2009; Rivas, et al., 2017). From the match score, we separated the pairing and unpairing portions of the match score calculation to derive scores for each nucleotide in each sequence, *PS*^*m*^(*i*) and *US*^*m*^(*i*), respectively (Equ.s 2-3), where the scores are calculated for the *i*^*th*^ nucleotide in the *m*^*th*^ sequence from a set. We generated two-dimension histograms of *US*^*m*^(*i*) as a function of *PS*^*m*^(*i*) for each sequence, where the histogram populations are for each nucleotide in the sequence (Fig. 6). We find that homologous sequences generally have a higher intensity closer to the diagonal compared to decoy sequences, and less intensity close to the origin (0,0). Therefore, the *PS*^*m*^*(i)* and *US*^*m*^*(i)* of a sequence provide a means to identify decoy sequences from homologous sequences.

**Figure 6.**
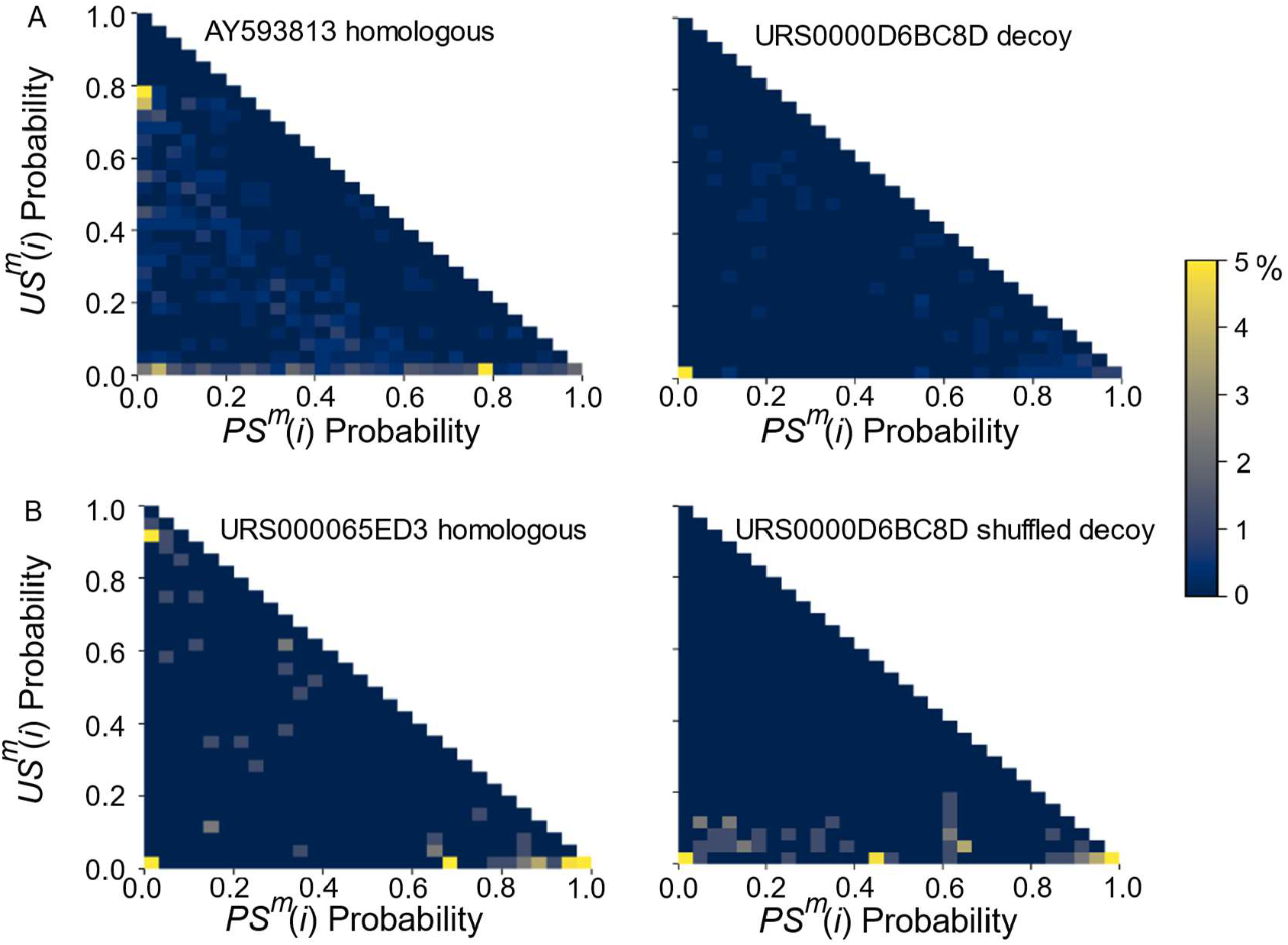
Heatmaps of *US*^*m*^(*i*) as a function of *PS*^*m*^(*i*) arrays for specified sequence sets calculated using TurboFold. (A) 8 IRES and 2 RAGATH sequences. AY593813 (left) represents a homologous IRES sequence and URS0000D6BC8D (right) represents a contamination sequence from the RAGATH family. (B) 15 RAGATH sequences and 1 shuffled decoy RAGATH sequence. URS000065ED3 (left) represents a homologous sequence, and URS0000D6BC8D decoy (right) represents shuffled decoy sequence.

### 3.4 AdaBoost identifies the decoy sequences in a group of homologous sequences

We applied a machine learning method, AdaBoost, to identify decoy sequences from homologous sequences (Pedregosa, et al., 2011). The method, called DecoyFinder, assesses each sequence, one at a time, using information from a TurboFold calculation and outputs the probability that a sequence is a decoy. Overall, the method is successful. Using all the features we discovered (explained below), DecoyFinder identifies 44.7% of decoys (sensitivity) with a rate of 5% for misidentifying homologs as decoys (1-specificity; blue line in Fig. 7). The threshold that provides this point on the ROC curve is 49.71% classification probability as decoy sequences.

**Figure 7.**
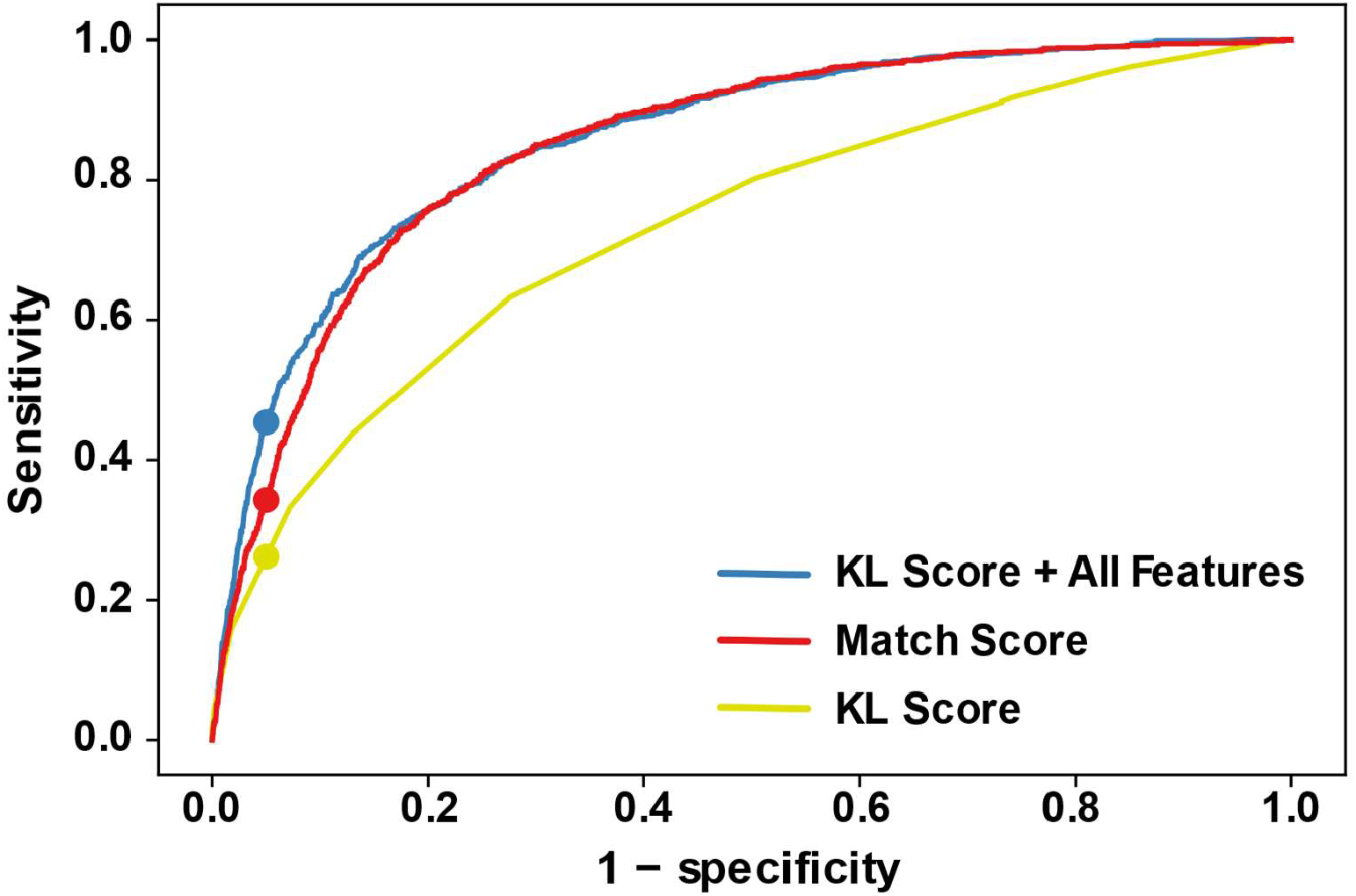
The ROC curve of the AdaBoost trained model. The ROC plots in blue, red, and yellow are obtained by using the KL score and all features, the *PS*^*m*^(*i*) and *US*^*m*^(*i*) histogram for the match scores, and KL score, respectively. The dots represent the sensitivity at which 1 – specificity = 0.05. The ROC curve is obtained by varying the AdaBoost model classification threshold for identifying contaminants in the test group. The area under the curve (AUC) of KL score and all feature is 0.857, the *PS*^*m*^(*i*) and *US*^*m*^(*i*) histogram for the match scores is 0.850 and KL score alone is 0.733.

DecoyFinder was trained and tested with separate families so that homologs did not appear in both sets (Lu, et al., 2009; Rivas, et al., 2012; Szikszai, et al., 2022). The details of the families are provided in the Methods section.

Using the histogram of *PS*^*m*^*(i)* as a function of *US*^*m*^*(i)*, the method is successful at identifying decoys (red line on Fig. 7). The PS and US scores, however, are a large 2-D histogram. To reduce this large histogram, the Kullback-Leibler divergence (KL divergence), which is a measure of difference between distributions (Kullback and Leibler, 1951), was used to summarize the information with a single score. We found that the use of a KL divergence summary was important when the other features were also integrated into the model. The full histogram apparently masked the other features and prevented them from improving the classification model. Utilizing the KL score in training model does decrease the accuracy of prediction (as quantified by area under the ROC curve, AUC; yellow line in Fig. 7) compared to the full 2-D array model when only the PS and US scores are used.

To increase the accuracy of identifying the decoy sequences, we added other features that help to distinguish the homologous sequences from the decoy sequences (see Methods). These include the Z-score of the folding free energy change, mean sequence Shannon entropy, the structural Shannon entropy for a single sequence calculation, the structural Shannon entropy after a TurboFold calculation and the difference in the two structural Shannon entropies. The Z-score of folding free energy represents how different the folding free energy change of a sequence is from the other sequences in the set. The mean alignment Shannon entropy reflects how well a sequence aligns with other sequences in the group. The structural Shannon entropy reflects the change in structures between a single sequence calculation and a TurboFold calculation including putative homologs. We hypothesize the decoy sequences have greater change because TurboFold will force them to fold into a structure that is closer to homologous sequences (Fig. 5). All these features can be obtained by post-processing after running TurboFold. The KL score in conjunction with the other features improves the model, with an increase sensitivity of prediction from about 0.3 to 0.45 at 0.95 specificity as compared to the *PS*^*m*^*(i)* and *US*^*m*^*(i)* histogram alone (red and blue curves in Fig. 7).

We also tested if the trained model identifies decoy sequences from other families and shuffled decoy sequences with different accuracies. The result shows that either test group with only decoy sequence or shuffled decoy sequence as contamination source have similar results (Supplementary Figure 2). It suggests that we are not overfitting our model to either type of decoy.

### 3.5 DecoyFinder implementation

DecoyFinder is easily used as an adjunct to TurboFold, and it is implemented in Python. DecoyFinder can directly take a TurboFold configuration file. TurboFold was modified to save the partition function files before TurboFold iterations and to output KL score, Z-score and difference in *SS(m)*. DecoyFinder can either run TurboFold or use TurboFold output files generated by a previously executed TurboFold calculation. After reading TurboFold output files, DecoyFinder calculates all the features and queries the trained machine learning model to identify if there is a potential decoy sequence. Finally, DecoyFinder outputs potential decoy sequences to suggest that they be removed from the TurboFold calculation (Fig. 1).

### 3.6 Paralogs represent a challenge to the model

Although we often use the term homologs for our sets of input sequences, following the practice in the field, we specifically mean orthologs, i.e. RNA genes that have a common ancestor and perform the same functions in separate species. We additionally tested our method using paralogs, which are RNA genes that arose from a duplication event and now have distinct functions. We used paralogs drawn from RNase P or RNase MRP and used the other family as the decoy. RNase P RNA and RNase MRP RNA share their catalytic Domain 1, but have a diverged Domain 2 (Li, et al., 2002; Piccinelli, et al., 2005).

We found that RNase MRP RNA as a decoy sequence in the RNase P RNA group (4 RNase P RNA and 1 RNase MRP RNA in each group and a total of 100 groups; Supplementary Table 5) was much more difficult to identify by our model compared to other test groups (AUC = 0.733; Figure 8, orange curve). RNase P RNA was even more difficult to identify as a contaminant in an RNase MRP RNA group, with an ROC curve such that the prediction result was close to randomly guessing (4 RNase MRP RNA and 1 RNase P RNA in each group and a total of 100 groups; AUC = 0.462; Figure 8, green curve). One possible explanation is that the heterogeneity in Domain 2 structures of RNase MRP RNA caused this difficulty. RNase P RNA and RNase MRP RNA share part of the structure in Domain 1, such as the P4 helix formed by CR-I and CR-IV. However, RNase P RNA have a high degree of conserved structure in Domain 2 across species, where, in contrast, RNase MRP RNA have divergent structures in Domain 2 (Li, et al., 2002; Piccinelli, et al., 2005). This makes it hard for DecoyFinder to distinguish MRP RNA as homologs and RNase P RNA as contaminants. On the other hand, in our test group where the homologs are RNase P RNA, DecoyFinder is less likely to identify RNase P RNA as decoys and still can identify RNase MRP correctly as decoys.

**Figure 8.**
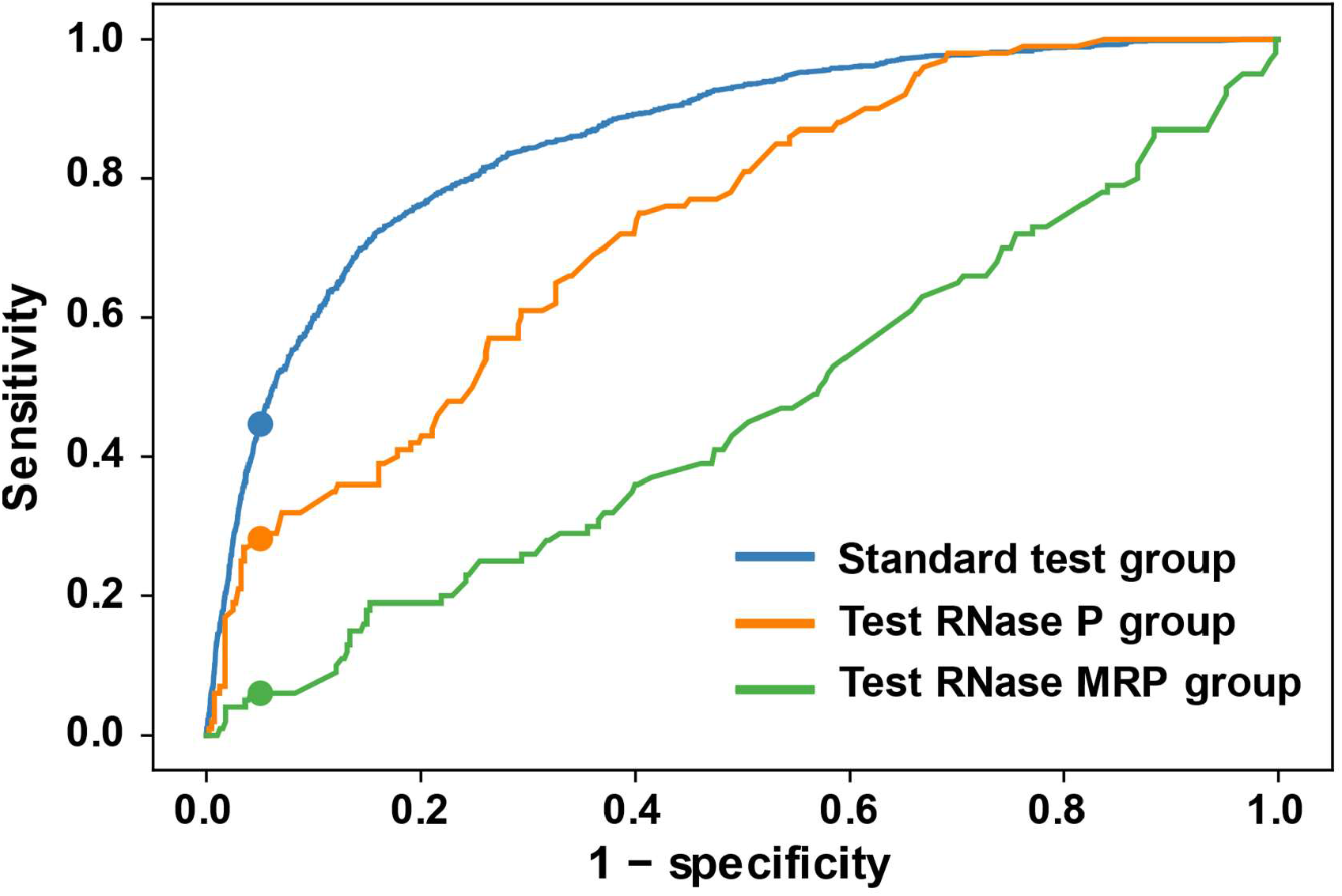
The ROC curve of DecoyFinder with RNase P RNA and RNase MRP RNA. The blue line represents the original test data across all families (Fig. 8). The orange line represents the test group whose homologous sequences are RNase P RNA and the decoy sequences are RNase MRP. The blue line represents a test group whose homologous sequences are RNase MRP RNA and the decoy sequences are RNase P RNA.

## 4 Discussion

### 4.1 TurboFold can tolerate decoy sequences in input homologous sequences

We find that the decoy sequences do not have a large effect on the accuracy of secondary structure prediction (Figure 3A-B). In fact, we observe that TurboFold tends to make a decoy sequence fold similarly to the homologs (Figure 5). These results demonstrate that TurboFold has some level of tolerance to decoy sequences contaminating the set of homologous input sequences (Tan, et al., 2017).

On the other hand, in spite of the loss in accuracy caused by contamination being small, the loss is statistically significant. Therefore, there is an advantage to identify and remove decoys to improve accuracy. Furthermore, identification of decoys will help to prevent contamination of databases of RNA families (Bateman, et al., 2011; Kalvari, et al., 2020; RNAcentral Consortium, 2021).

### 4.2 DecoyFinder is Unique

To our knowledge, there is currently no software that provides a function like DecoyFinder, but Infernal is the tool of choice once a well-established RNA family model is developed (Altschul, et al., 1990; Nawrocki and Eddy, 2013). For instance, Infernal can take a structurally annotated alignment of an RNA family to search for homologous sequences (Nawrocki and Eddy, 2013). In contrast to Infernal, which is trained on each family using a well-established alignment, DecoyFinder needs only the small set of putative homolog sequences and does not rely on having an established structure for the family because all features are directly derived from TurboFold predictions. DecoyFinder will be used early in the process of comparative analysis to prevent downstream contamination of the RNA family. In turn, this early curation of a nascent family can improve the training of Infernal.

### 4.3 The features of DecoyFinder might contribute to improving the accuracy of homolog search methods

Several software tools are available to identify RNA sequence homologs early in the process of determining a conserved structure. Rsearch, RNAlien, RNAcmap, and rMSA provide tools for scanning sequence databases for homologs (Eggenhofer, et al., 2016; Klein and Eddy, 2003; Zhang, et al., 2022; Zhang, et al., 2021). Additionally, RNAz, Multifind, and CMFinder can identify novel ncRNAs in sequence data (Fu, et al., 2015; Gruber, et al., 2010; Torarinsson, et al., 2008). These tools are prone to discovery of false positives, and DecoyFinder might prove useful as a method that can provide additional data in the classification of sequences. Additionally, the features that we use in the identification of contaminating sequences might prove useful in the further development of other software tools.

## 5 Acknowledgements

This work was supported by NIH grant R35 GM145283 to D.H.M. The University of Rochester Center for Integrated Research Computing provided computational resources.

## Notes

### Competing Interest Statement

The authors have declared no competing interest.

