## Supplemental Figures and Tables for "DecoyFinder: Identification of Contaminants in Sets of Homologous RNA Sequences"

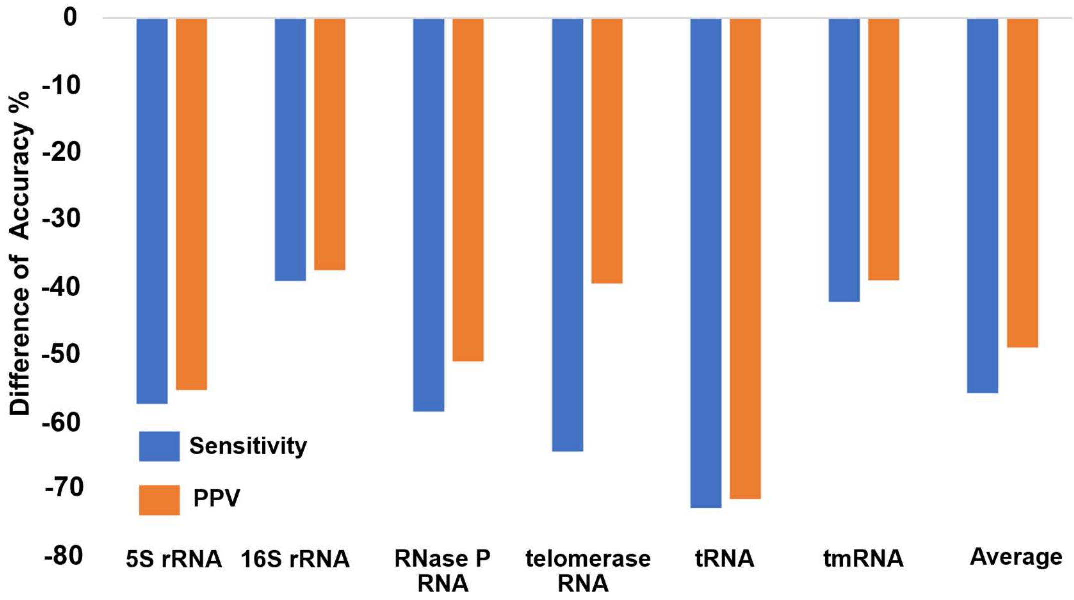


**Supplementary Figure 1**. Secondary structure prediction accuracy difference between shuffled decoy sequences and natural RNA sequences. Structures were predicted using maximum expected accuracy prediction (Lu, et al., 2009). For each RNA family, 100 sequences were chosen at random to generate a shuffled decoy. Sensitivity and PPV were calculated by comparing the predicted structure of shuffled decoy sequences with known structure of original sequence using the Scorer program in RNAstructure. The average change of sensitivity was -55.98% and the average change in PPV was -49.14%

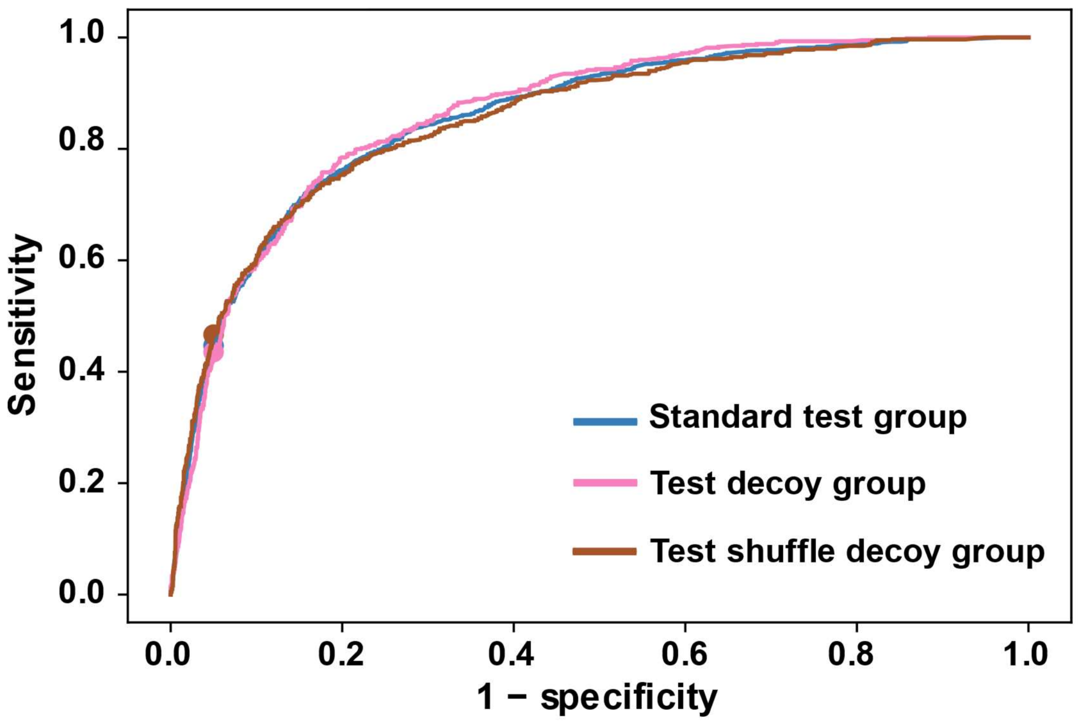


**Supplementary Figure 2**. ROC curves of the AdaBoost trained model for the standard test group with both types of decoys (blue), the test group with decoys from other families (pink), and the test group with shuffled decoys (brown). In both the decoy group and the shuffled decoy group, there is only one type of decoy.

**Supplementary Table 1.** The training set for the machine learning model. The Rfam families are 5S rRNA, internal ribosome entry site (IRES), malK-I RNA, RAGATH-18 RNA, RNase P RNA, skipping rope RNA, tRNA and U3 RNA. There are 900 sets of decoys from other families, 900 sets of shuffled decoys and 300 sets that do not include decoys. The sequences were randomly selected without replacement, therefore if a sequence has been chosen for one type of set (in a set of another family, shuffled decoy, or non-decoy) it will not be selected again. The selection of homologous and decoy families was random, but for each of the two types of decoys there are a total of 300 sets containing 1 decoy, 300 sets with 2 decoys and 300 sets with 3 decoys. The number of homologous sequences in all sets are randomly selected, ranging from 5 to 20.

| Number of sets | Sequence family in set | Decoys in set | Decoy type (shuffle/other family/no decoy) | Sequence source | Sequences in set |
|---|---|---|---|---|---|
| 13 | 5S rRNA | IRES | other family | Rfam | 170 |
| 9 | 5S rRNA | RAGATH | other family | Rfam | 139 |
| 11 | 5S rRNA | RNase P | other family | Rfam | 168 |
| 14 | 5S rRNA | U3 | other family | Rfam | 212 |
| 11 | 5S rRNA | malK | other family | Rfam | 180 |
| 14 | 5S rRNA | skipping rope RNA | other family | Rfam | 196 |
| 29 | 5S rRNA | tRNA | other family | Rfam | 417 |
| 11 | IRES | 5S | other family | Rfam | 154 |
| 15 | IRES | RAGATH | other family | Rfam | 203 |
| 24 | IRES | RNase P | other family | Rfam | 323 |
| 18 | IRES | U3 | other family | Rfam | 275 |
| 14 | IRES | malK | other family | Rfam | 198 |
| 17 | IRES | skipping rope RNA | other family | Rfam | 240 |
| 14 | IRES | tRNA | other family | Rfam | 161 |
| 20 | RAGATH | 5S | other family | Rfam | 307 |
| 19 | RAGATH | IRES | other family | Rfam | 276 |
| 13 | RAGATH | RNase P | other family | Rfam | 197 |
| 13 | RAGATH | U3 | other family | Rfam | 202 |
| 12 | RAGATH | malK | other family | Rfam | 190 |
| 14 | RAGATH | skipping rope RNA | other family | Rfam | 164 |
| 21 | RAGATH | tRNA | other family | Rfam | 287 |
| 13 | RNase P | 5S | other family | Rfam | 184 |
| 18 | RNase P | IRES | other family | Rfam | 247 |
| 14 | RNase P | RAGATH | other family | Rfam | 201 |
| 15 | RNase P | U3 | other family | Rfam | 198 |
| 15 | RNase P | malK | other family | Rfam | 250 |
| 11 | RNase P | skipping rope RNA | other family | Rfam | 166 |
| 16 | RNase P | tRNA | other family | Rfam | 218 |
| 15 | U3 | 5S | other family | Rfam | 206 |

| | | | | | |
|---|---|---|---|---|---|
| 15 | U3 | IRES | other family | Rfam | 225 |
| 9 | U3 | RAGATH | other family | Rfam | 122 |
| 14 | U3 | RNase P | other family | Rfam | 198 |
| 25 | U3 | malK | other family | Rfam | 347 |
| 18 | U3 | skipping rope RNA | other family | Rfam | 279 |
| 22 | U3 | tRNA | other family | Rfam | 318 |
| 16 | malK | 5S | other family | Rfam | 217 |
| 17 | malK | IRES | other family | Rfam | 247 |
| 16 | malK | RAGATH | other family | Rfam | 265 |
| 18 | malK | RNase P | other family | Rfam | 249 |
| 24 | malK | U3 | other family | Rfam | 317 |
| 12 | malK | skipping rope RNA | other family | Rfam | 184 |
| 13 | malK | tRNA | other family | Rfam | 195 |
| 25 | skipping rope RNA | 5S | other family | Rfam | 373 |
| 8 | skipping rope RNA | IRES | other family | Rfam | 131 |
| 13 | skipping rope RNA | RAGATH | other family | Rfam | 195 |
| 15 | skipping rope RNA | RNase P | other family | Rfam | 202 |
| 21 | skipping rope RNA | U3 | other family | Rfam | 287 |
| 24 | skipping rope RNA | malK | other family | Rfam | 338 |
| 9 | skipping rope RNA | tRNA | other family | Rfam | 108 |
| 18 | tRNA | 5S | other family | Rfam | 262 |
| 19 | tRNA | IRES | other family | Rfam | 289 |
| 14 | tRNA | RAGATH | other family | Rfam | 212 |
| 19 | tRNA | RNase P | other family | Rfam | 234 |
| 14 | tRNA | U3 | other family | Rfam | 192 |
| 22 | tRNA | malK | other family | Rfam | 326 |
| 17 | tRNA | skipping rope RNA | other family | Rfam | 229 |
| 170 | 5S rRNA | 5S rRna | Shuffle | Rfam | 2455 |
| 24 | IRES | IRES | Shuffle | Rfam | 349 |
| 189 | RAGATH | RAGATH | Shuffle | Rfam | 2797 |
| 29 | RNase P | RNase P | Shuffle | Rfam | 412 |
| 21 | U3 | U4 | Shuffle | Rfam | 316 |
| 82 | malK | malK | Shuffle | Rfam | 1123 |
| 204 | skipping rope RNA | skipping rope RNA | Shuffle | Rfam | 2825 |
| 181 | tRNA | tRNA | Shuffle | Rfam | 2670 |
| 52 | 5S rRNA | no decoy | no decoy | Rfam | 691 |
| 8 | IRES | no decoy | no decoy | Rfam | 97 |
| 69 | RAGATH | no decoy | no decoy | Rfam | 888 |

| | | | | | |
|---|---|---|---|---|---|
| 9 | RNase P | no decoy | no decoy | Rfam | 126 |
| 7 | U3 | no decoy | no decoy | Rfam | 89 |
| 28 | malK | no decoy | no decoy | Rfam | 324 |
| 57 | skipping rope RNA | no decoy | no decoy | Rfam | 656 |
| 70 | tRNA | no decoy | no decoy | Rfam | 950 |

**Supplementary Table 2.** The testing sets for DecoyFinder. The Rfam families are pemK RNA motif, small nucleolar RNA SNORD116, U4 spliceosomal RNA, U5 spliceosomal RNA and Y RNA.  There are 300 sets of decoys from other families, 300 sets of shuffled decoys and 100 sets that contain no decoy. The sequences were randomly selected without replacement, therefore if a sequence has been chosen for one type of set (in a set of another family, shuffled decoy, or non-decoy) it will not be selected again. The selection of homologous and decoy families was random, but for each of the two decoy types there are a total of 100 sets containing 1 decoy, 100 sets with 2 decoys and 100 sets with 3 decoys. The number of homologous sequences in all sets are randomly selected, ranging from 5 to 20.

| Number of sets | Sequence family in set | Decoys in set | Decoy type (shuffle/other family/non) | Sequence source | Sequence in set |
|---|---|---|---|---|---|
| 18 | SNORD116 | Y RNA | other family | Rfam | 214 |
| 18 | SNORD117 | U4 | other family | Rfam | 243 |
| 12 | SNORD118 | U5 | other family | Rfam | 132 |
| 11 | SNORD119 | pemK RNA motify | other family | Rfam | 127 |
| 15 | U4 | Y RNA | other family | Rfam | 217 |
| 15 | U4 | SNORD116 | other family | Rfam | 238 |
| 15 | U4 | U5 | other family | Rfam | 210 |
| 17 | U4 | pemK RNA motify | other family | Rfam | 234 |
| 15 | U5 | Y RNA | other family | Rfam | 231 |
| 15 | U5 | SNORD116 | other family | Rfam | 197 |
| 10 | U5 | U4 | other family | Rfam | 160 |
| 13 | U5 | pemK RNA motify | other family | Rfam | 168 |
| 21 | Y RNA | SNORD116 | other family | Rfam | 312 |
| 23 | Y RNA | U4 | other family | Rfam | 333 |
| 13 | Y RNA | U5 | other family | Rfam | 203 |
| 12 | Y RNA | pemK RNA motify | other family | Rfam | 172 |
| 18 | pemK RNA motify | Y RNA | other family | Rfam | 276 |
| 17 | pemK RNA motify | SNORD116 | other family | Rfam | 252 |
| 11 | pemK RNA motify | U4 | other family | Rfam | 150 |
| 11 | pemK RNA motify | U5 | other family | Rfam | 160 |
| 14 | SNORD116 | SNORD117 | Shuffle | Rfam | 198 |
| 44 | U4 | U4 | Shuffle | Rfam | 614 |
| 41 | U5 | U5 | Shuffle | Rfam | 664 |
| 23 | Y RNA | Y RNA | Shuffle | Rfam | 372 |
| 178 | pemK RNA | pemK RNA | Shuffle | Rfam | 2597 |

| | | | | | |
|---|---|---|---|---|---|
| | motify | motify | | | |
| 4 | SNORD116 | non | non | Rfam | 54 |
| 17 | U4 | non | non | Rfam | 173 |
| 13 | U5 | non | non | Rfam | 186 |
| 8 | Y RNA | non | non | Rfam | 105 |
| 58 | pemK RNA motify | non | non | Rfam | 769 |

**Supplementary Table 3.** The testing set for TurboFold's robustness with contamination present. The RNAStralign families used are 5S rRNA, 16S rRNA, RNase P RNA, telomerase RNA, tRNA and tmRNA. There are 700 sets of decoys taken from other families and 700 sets of shuffled decoys. Each set contains 4 homologous sequences and 1 decoy sequence. The sequences were randomly selected without replacement, therefore if a sequence has been chosen for one type of set (in a set of another family, shuffled decoy, or non-decoy) it will not be selected again.

| Number of sets | Sequence family in set | Decoys in set | Decoy type (shuffle/other family/non) | Sequence source |
|---|---|---|---|---|
| 15 | 5S rRNA | RNase P | other family | RNAStralign |
| 18 | 5S rRNA | SRP | other family | RNAStralign |
| 13 | 5S rRNA | 16S rRNA | other family | RNAStralign |
| 30 | 5S rRNA | tRNA | other family | RNAStralign |
| 12 | 5S rRNA | telomerase RNA | other family | RNAStralign |
| 12 | 5S rRNA | tmRNA | other family | RNAStralign |
| 13 | 16S rRNA | RNase P | other family | RNAStralign |
| 17 | 16S rRNA | SRP | other family | RNAStralign |
| 20 | 16S rRNA | 5S rRna | other family | RNAStralign |
| 23 | 16S rRNA | tRNA | other family | RNAStralign |
| 11 | 16S rRNA | telomerase RNA | other family | RNAStralign |
| 16 | 16S rRNA | tmRNA | other family | RNAStralign |
| 14 | RNase P | 16S rRNA | other family | RNAStralign |
| 20 | RNase P | 5S rRNA | other family | RNAStralign |
| 22 | RNase P | SRP | other family | RNAStralign |
| 9 | RNase P | tRNA | other family | RNAStralign |
| 20 | RNase P | telomerase RNA | other family | RNAStralign |
| 15 | RNase P | tmRNA | other family | RNAStralign |
| 18 | SRP | 16S rRNA | other family | RNAStralign |
| 8 | SRP | 5S rRNA | other family | RNAStralign |
| 13 | SRP | RNase P | other family | RNAStralign |
| 23 | SRP | tRNA | other family | RNAStralign |
| 23 | SRP | telomerase RNA | other family | RNAStralign |
| 15 | SRP | tmRNA | other family | RNAStralign |
| 16 | telomerase RNA | 16S rRNA | other family | RNAStralign |
| 22 | telomerase RNA | 5S rRNA | other family | RNAStralign |
| 16 | telomerase RNA | RNase P | other family | RNAStralign |
| 15 | telomerase RNA | SRP | other family | RNAStralign |
| 14 | telomerase | tRNA | other family | RNAStralign |

| | | | | |
|---|---|---|---|---|
| | RNA | | | |
| 17 | telomerase RNA | tmRNA | other family | RNAStralign |
| 16 | tmRNA | 16S rRNA | other family | RNAStralign |
| 17 | tmRNA | 5S rRNA | other family | RNAStralign |
| 21 | tmRNA | RNase P | other family | RNAStralign |
| 9 | tmRNA | SRP | other family | RNAStralign |
| 18 | tmRNA | tRNA | other family | RNAStralign |
| 19 | tmRNA | telomerase RNA | other family | RNAStralign |
| 23 | tRNA | 16S rRNA | other family | RNAStralign |
| 13 | tRNA | 5S rRNA | other family | RNAStralign |
| 16 | tRNA | RNase P | other family | RNAStralign |
| 11 | tRNA | SRP | other family | RNAStralign |
| 17 | tRNA | telomerase RNA | other family | RNAStralign |
| 20 | tRNA | tmRNA | other family | RNAStralign |
| 100 | 16S rRNA | 16S rRNA | Shuffle | RNAStralign |
| 100 | 5S rRNA | 5S rRNA | Shuffle | RNAStralign |
| 100 | RNase P | RNase P | Shuffle | RNAStralign |
| 100 | SRP | SRP | Shuffle | RNAStralign |
| 100 | telomerase RNA | telomerase RNA | Shuffle | RNAStralign |
| 100 | tmRNA | tmRNA | Shuffle | RNAStralign |
| 100 | tRNA | tRNA | Shuffle | RNAStralign |

**Supplementary Table 4.** The RNA families' pairwise identity. The pairwise identity was determined by 30 randomly selected sequences from the TurboFold alignment.

| Sequence family | Mean sequence identity | Usage |
|---|---|---|
| 5S rRNA | 56.9 | Shuffle sequence accuracy test |
| 16S rRNA | 79.4 | Shuffle sequence accuracy test |
| RNase P RNA | 47.1 | Shuffle sequence accuracy test |
| telomerase RNA | 62.9 | Shuffle sequence accuracy test |
| tRNA | 50.6 | Shuffle sequence accuracy test |
| tmRNA | 49.7 | Shuffle sequence accuracy test |
| 5S rRNA | 56.4 | Model Training |
| internal ribosome entry site (IRES) | 82.7 | Model Training |
| malK-I RNA | 58.6 | Model Training |
| RAGATH-18 RNA | 60.6 | Model Training |
| RNase P RNA | 47.9 | Model Training |
| skipping rope RNA | 53.8 | Model Training |
| tRNA | 42.2 | Model Training |
| U3 RNA | 65.9 | Model Training |
| pemK RNA motif | 65 | Model Testing |
| small nucleolar RNA SNORD116 | 82.2 | Model Testing |
| U4 spliceosomal RNA | 59.8 | Model Testing |
| U5 spliceosomal RNA | 54 | Model Testing |
| Y RNA | 70.6 | Model Testing |

**Supplementary Table 5.** The RNase P RNA and RNase MRP RNA testing sets for DecoyFinder each contain 100 groups, with one group being homologous and the other being the decoy. The sequences were randomly selected without replacement; therefore, if a sequence has been chosen for one type of set (in a set of another family or non-decoy), it will not be selected again. The selection of homologous and decoy families was random, and each test set contains four homologous sequences and one decoy sequence from the other family.

| Number of sets | Sequence family in set | Decoys in set | Decoy type (shuffle/other family/no decoy) | Sequence source | Sequences in set |
|---|---|---|---|---|---|
| 100 | RNase P | RNase MRP | other family | Rfam | 500 |
| 100 | RNase MRP | RNase P | other family | Rfam | 500 |
